# Amino acid availability determines plant immune homeostasis in the rhizosphere microbiome

**DOI:** 10.1101/2021.12.20.473424

**Authors:** Yang Liu, Jiatong Han, Andrew J. Wilson, Lucy O’Sullivan, Cara H. Haney

## Abstract

Microbes possess conserved microbe-associated molecular patterns (MAMPs) such as flagellin that are recognized by plant receptors to induce immunity. Despite containing the same MAMPs as pathogens, commensals thrive in the plant rhizosphere microbiome indicating they must suppress or evade host immunity. The beneficial bacteria *Pseudomonas capeferrum* WCS358 can suppress *Arabidopsis* root immunity via acidification by secreting gluconic acid. While gluconic acid is sufficient to suppress immunity, we found that it is not necessary in a second beneficial strain *Pseudomonas simiae* WCS417, which produces more gluconic acid than WCS358. To uncover mechanisms that contribute to the suppression of *Arabidopsis* immunity, we performed a forward genetic screen in EMS-mutagenized *P. simiae* WCS417 using a flagellin-inducible *CYP71A12_pro_:GUS* reporter as an *Arabidopsis* immune readout. We identified a mutant that cannot suppress flagellin-elicited *CYP71A12_pro_:GUS* expression or acidify the rhizosphere. Next generation sequencing revealed a mutation in the catabolic site of an ornithine carbamoyltransferase *argF*, which is required for arginine biosynthesis. The mutant could be complemented by expression of *argF* from a plasmid, and a Δ*argF* mutant could not suppress immunity. Fungal pathogens can use alkalization through production of ammonia and glutamate, the arginine biosynthetic precursors, to promote their own growth and virulence. Therefore, we hypothesized that the biosynthesis of specific amino acids may be necessary to reduce levels of ammonia and glutamate to prevent rhizosphere alkalization and bacterial overgrowth. Genetically blocking arginine, glutamine, or proline biosynthesis, or by adding corresponding exogenous amino acids, resulted in rhizosphere alkalization. Interestingly, exogenous amino acids caused bacterial overgrowth in a gluconic acid-deficient mutants. Our findings show that bacterial amino acid biosynthesis contributes to acidification by preventing accumulation of glutamate precursors and the resulting alkalization. Collectively this work shows that by regulating nutrient availability, plants have the potential to regulate their immune homeostasis in the rhizosphere microbiome.

## Introduction

A myriad of microorganisms known as microbiota dwell in the plant rhizosphere. The interactions between the plant and its associated microbiota actively influence the fitness of one another [1]. Both pathogenic and mutualistic microbes form associations with plant. To protect themselves from pathogens, the plant immune system diligently monitors the environment to prevent pathogenic invasion. To do so, plants possess pattern recognition receptors (PRRs) that can specifically sense microbe-associated molecular patterns (MAMPs) that are evolutionarily conserved across diverse microbes. Perception of MAMPs triggers pattern-triggered immunity (PTI), which includes a reactive oxygen species burst, calcium influx and defense gene expression [2]. As both pathogens and commensals contain MAMPs that can be recognized by the plant innate immune system, both must supress, or evade immunity in order to successfully colonize. While mechanisms of immunity suppression by pathogenic bacteria are well-established, how commensal microbes evade or suppress plant immunity to promote their own fitness is poorly understood.

Some successful pathogens can suppress PTI by injecting effector proteins into the plant cytosol via type three secretion systems (T3SS) to suppress PTI [3]. In addition to injecting effectors, pathogenic bacteria can manipulate phytohormones to suppress host immunity. The pathogenic bacterium *Pseudomonas syringae* pv tomato DC3000 (*Pto*) can secrete the phytotoxin Coronatine (COR) to mimic the active form of jasmonic acid (JA), JA-lIe, promoting JA-dependent defense. Since the JA and salicylic acid (SA) pathways antagonize each other, inducing JA signaling by COR suppresses SA, which is critical for resistance against *Pto* [4]. Lastly, instead of suppressing host immunity, pathogenic microbes can hide their MAMPs to prevent recognition by PRRs. For instance, *P. syringae* can secrete AprA, an extracellular alkaline protease that can degrade flagellin monomers, thereby avoiding immune recognition [5].

Although commensals also have MAMPs that have the potential to induce PTI, they do not induce immune responses, suggesting that commensals can suppress or evade the plant immune system [6]. A growth promoting and biocontrol strain *Pseudomonas* sp. WCS365 can evade host immunity by fine tuning biofilm formation [7]. *Pseudomonas capeferrum* WCS358 can suppress root local immunity by secreting organic acids to lower the pH of the rhizosphere [8]. *Dyella japonica* suppresses root immunity through a Type II secretion-dependent mechanism without affecting rhizosphere pH [9]. Collectively these findings indicate that rhizosphere commensal possess diverse mechanisms to modulate host immunity.

The beneficial root-associated bacterial strain, *Pseudomonas simiae* WCS417, was previously shown to suppress root immunity [8,10]. It lowers the pH of the rhizosphere to a greater extent than *P. capeferrum* WCS358 and produces more gluconic acid [8]; however, we found that deletion of *pqqF*, which results in loss of gluconic acid biosynthesis and immunity suppression in *P. capeferrum* WCS358 [8], does not impair the ability of WCS417 to suppress rhizosphere immunity. Here we describe a forward genetic screen which identified a novel mechanism of root immunity suppression in WCS417, that bacteria employ specific amino acids biosynthesis to prevent rhizosphere alkalization and suppress immunity.

## Results

### Acidification is sufficient but not necessary for *Arabidopsis* immunity suppression by *P. simiae* WCS417

Gluconic acid biosynthesis via *pqqF* is necessary for *Arabidopsis* rhizosphere immunity suppression in *P. capeferrum* WCS358 and *P. aeruginosa* PAO1 as measured by expression of a PTI-inducible reporter *CYP71A12_pro_:GUS* expression [8]. *CYP71A12* is involved in biosynthesis of the antimicrobial camalexin and is induced in the root elongation zone or maturation zone upon sensing MAMPs such as flg22 or chitin [10]. As a result, induction of *CYP71A12_pro_:GUS* reports the activation of PTI. *P. simiae* WCS417 produces more gluconic acid and lowers the pH of the rhizosphere to a greater extent than WCS358 suggesting WCS417 *pqqF* is also required for rhizosphere immunity suppression[8]. However, we found that while a clean deletion of *pqqF* in *P. simiae* WCS417 resulted in a significant increase in the pH of seedling exudates (Figure 1A), the WCS417 Δ*pqqF* mutant can still suppress flg22-triggered expression of *CYP71A12pro:GUS* (Figure 1B). In contrast, disruption of *pqqF* from *Pseudomonas aeruginosa* PAO1 resulted in a less dramatic increase in the pH of seedling exudates (Figure 1A) and has been shown necessary for host immunity suppression [8]. These data suggest that WCS417 possesses additional mechanisms to suppress host immunity.

**Figure 1.**
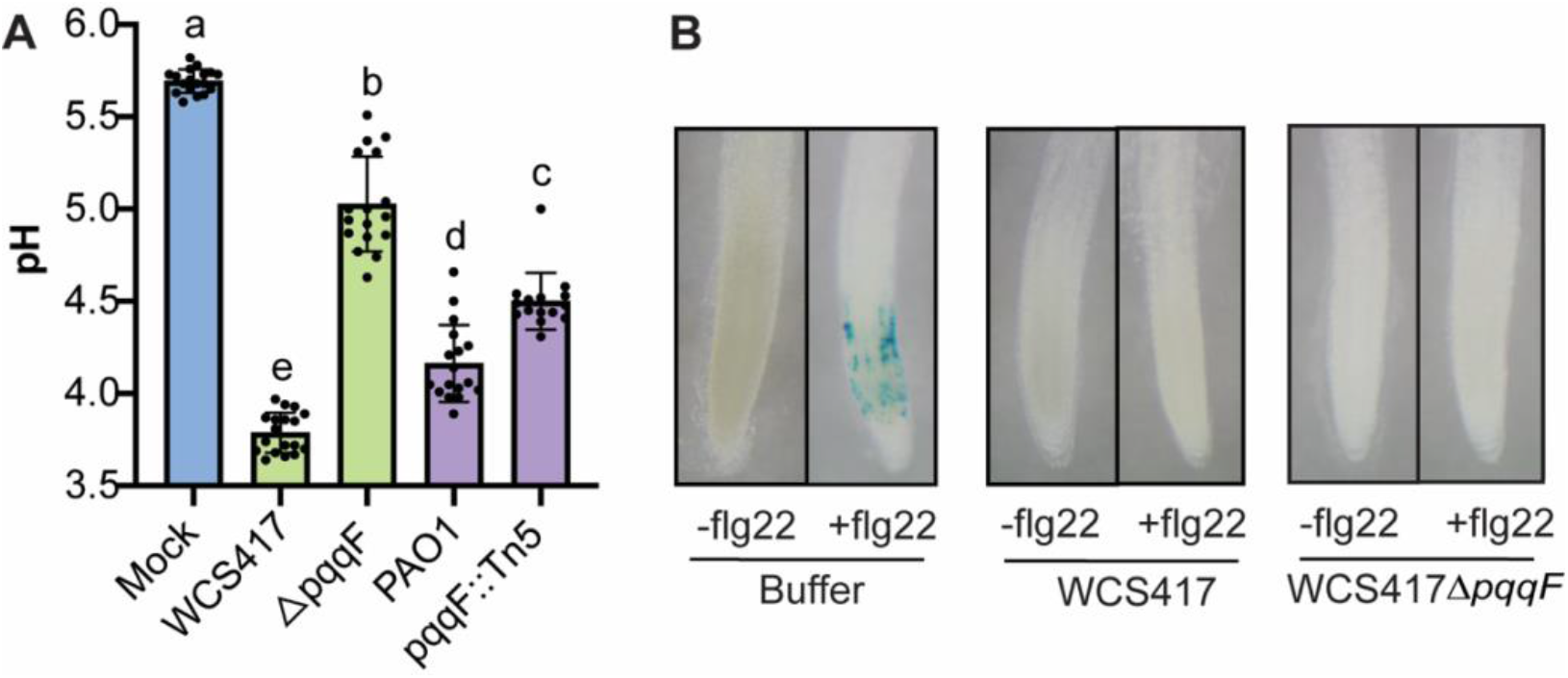
*pqqF* is not required for host immunity suppression in the *P. simiae* WCS417. (A) Deletion or disruption of *pqqF* results in a pH increase in WCS417 and *P. aeruginosa* PAO1. Individual data are the result of a single well of seedling exudates. Statistics were calculated by using one-way ANOVA and Tukey’s HSD. Error bars represent mean +/− SD and letters indicate differences at p<0.05. (B) While the Δ*pqqF* mutant increased pH, it can still suppress flg22-induced *CYP71A12_pro_:GUS* expression. All the experiments were independently repeated at least 3 times.

### *P. simiae* WCS417 *argF* is required for rhizosphere acidification and immunity suppression

To identify additional genes that are necessary for host immunity suppression, we generated an EMS-mutagenized library of *P. simiae* WCS417 and screened for mutants that were unable to suppress flg22-mediated induction of the *CYP71A12_pro_:GUS* reporter. We screened 960 EMS-mutagenized colonies of WCS417 in duplicate for their ability to suppress flg22-induced expression of the *CYP71A12_pro_:GUS* reporter. A single mutant named 10E10 was identified that cannot suppress flg22-induced immunity (Figure 2A). We found that 10E10 completely failed to reduce the pH of seedling exudates suggesting that it might contribute to immunity suppression through rhizosphere acidification (Figure 2B).

**Figure 2.**
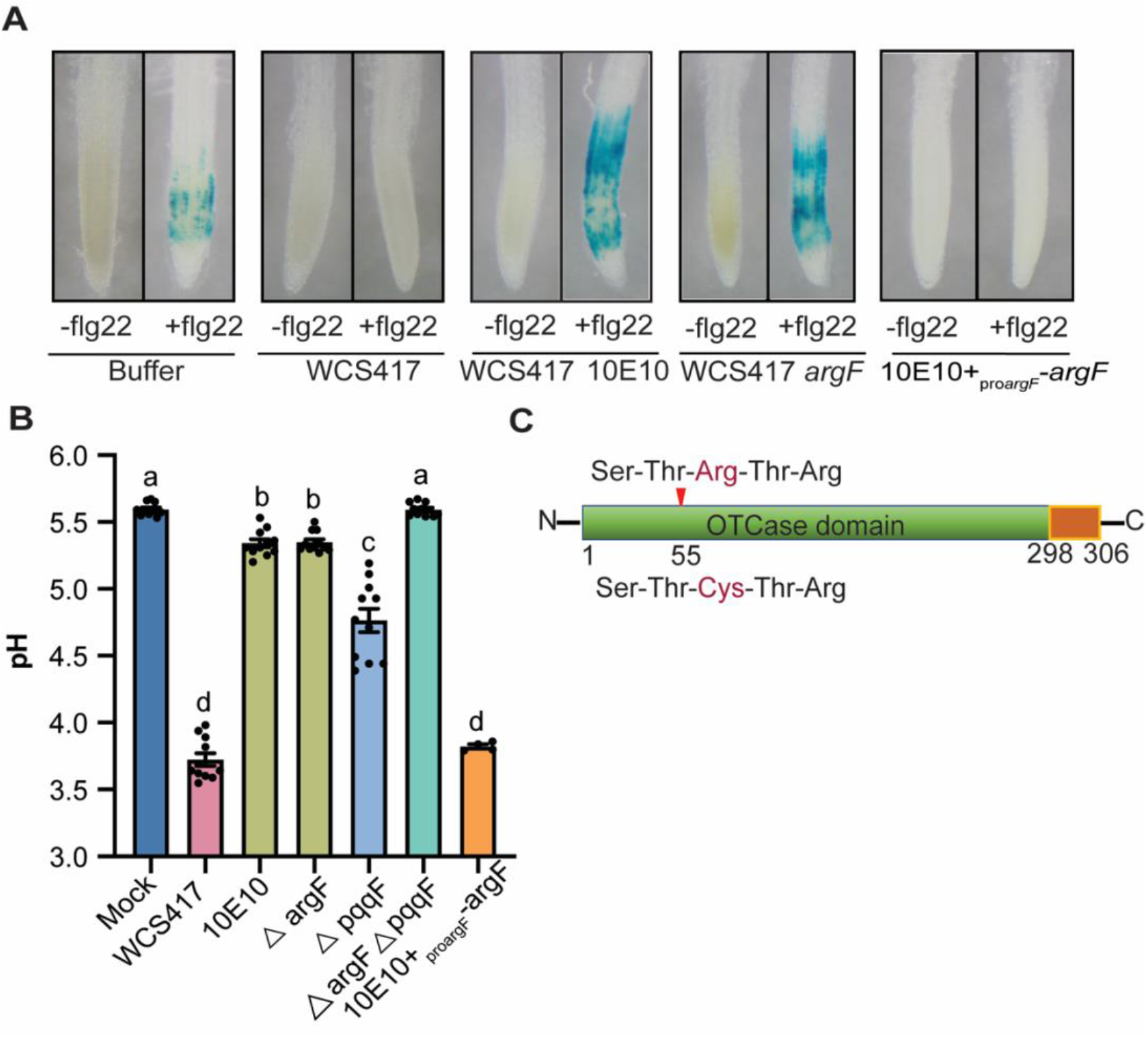
An EMS screen identified a loss of function mutant in Δ*argF* that cannot suppress *Arabidopsis* immunity or lower rhizosphere pH. (A) A mutant, 10E10 was identified from an EMS screen for *P. simiae* mutants that cannot supress flg22-triggered induction of *CYP71A12_pro_:GUS* expression or (B) acidify the rhizosphere. An Δ*argF* mutant phenocopied the inability of the 10E10 mutant to suppress immunity suppression and acidification, and expression of *_proargF_-argF* on a plasmid restored the immunity suppression and acidification of the 10E10 mutant. (C)The mutation in the *argF* gene in the 10E10 mutant is within its catalytic site resulting in a predicted loss of function mutation. All the experiments were independently repeated at least 3 times. Statistics were calculated by using one-way ANOVA and Tukey’s HSD. Error bars represent mean +/− SD and letters indicate differences at p<0.05

Because we only identified one mutant from the screen, we wondered if mutations in WCS417 that result in a loss of immunity suppression are rare, or if our screen was not saturating. To test this, we determined whether we could identify an allele of *pqqF* in our screen. Gluconic acid has previously been shown be required for zinc solubilization and so we tested whether we could identify a mutant unable to solubilize zinc. By growing bacteria on zinc phosphate media [11], only the strains that can solubilize zinc phosphate will produce a clear halo on the plate. We found a single mutant 4E4 that cannot solubilize zinc phosphate (Figure S1). The genome of 4E4 was sequenced and we identified mutations in *pqqF* (Table S2). However, 4E4 can still suppress flg22-triggered *CYP71A12_pro_:GUS* expression, which is consistent with our finding that Δ*pqqF* WCS417 can still suppress host immunity. As our screen identified a mutation in *pqqF*, this suggests that mutants that fail to suppress root immunity might be rare in *P. simiae* WCS417.

We sequenced the genome of the *P. simiae* WCS417 mutant 10E10 and identified 35 non-synonymous mutations with respect to the parental WCS417 strain (Table S1). To narrow down candidate genes, we made use of a PAO1 transposon insertion library [12] and tested transposon insertion mutants in genes from the mapping list that we hypothesized were most likely to contribute to immunity suppression. We found that an insertion in *argF* uniquely impaired the ability of PAO1 to acidify seedling exudates (Figure 2A, Figure S2A). *ArgF* encodes ornithine carbamoyltransferase, which is involved in arginine biosynthesis, converting L-ornithine to L-citrulline [13,14]. The mutation in *argF* in *P. simiae* WCS417 is predicted to affect its catalytic site Ser-Thr-Arg-Thr-Arg, where a C to T change is predicted to convert the third arginine to cysteine (Figure 2C). Thus, we hypothesized that a loss of function of *argF* likely underlines the WCS417-mediated immunity suppression.

We tested whether a loss of *P. simiae* WCS417 *argF* could explain the inability of the 10E10 mutant to grow in the rhizosphere and suppress immunity. First, we confirmed the C to T mutation in the *argF* catalytic site in the 10E10 mutant by PCR. We complemented the 10E10 mutant with *argF* expressed by its native promoter in the PBBR1MCS-5 plasmid and found that *_proargF_-argF* rescued the 10E10 mutant and the complemented strain suppressed flg22-triggered *CYP71A12_pro_:GUS* expression, resulting in acidification of seedling exudates to a similar degree as wildtype WCS417 (Figure 2A, B). We then made a clean deletion of *argF* in WCS417 and found that the *ΔargF* mutant phenocopied the inability of the 10E10 mutant to suppress flg22-triggered *CYP71A12_pro_:GUS* expression and could not acidify seedling exudates (Figure 2A, B). These results illustrate that a loss of function of *argF* underlies the inability to suppress immunity by the 10E10 mutant.

### Conversion of ammonia and glutamate to arginine, proline, and glutamine, is necessary for bacterial growth and to avoid rhizosphere alkalization

A loss of function mutation in *argF* may result in accumulation of arginine precursors including L-ornithine and TCA intermediates and a defect in generating protons, L-citrulline, arginine (Figure 3A). Thus, it is possible that downstream metabolites from *argF*, or feedback inhibition on an upstream pathway, underly the inability of immunity suppression of 10E10. Fungal pathogens can produce glutamate, which is a biosynthetic precursor of arginine, to hijack plant metabolism, increase apoplastic pH, and promote their own virulence [15,16]. As a result, we hypothesized that a loss of *argF* may result in accumulation of glutamate and ammonia, resulting in alkalization of the rhizosphere.

**Figure 3.**
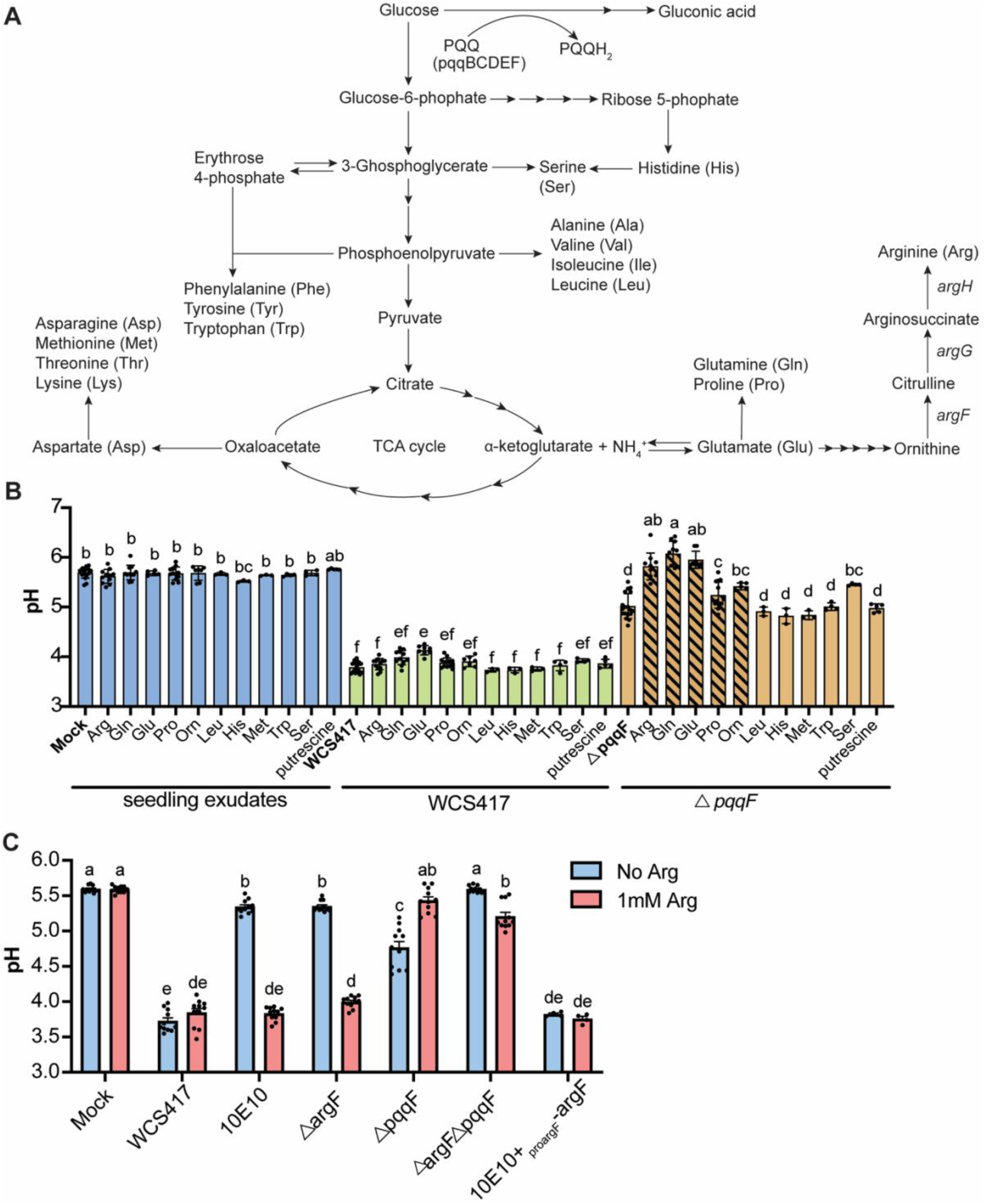
Glutamate biosynthesis pathway contributes to rhizosphere acidification, but the effect is masked by gluconic acid biosynthesis. (A) Glutamate and gluconic acid biosynthesis pathways in *Pseudomonas*. (B) Blocking glutamate biosynthesis by providing arginine, proline, glutamine, and glutamate (striped bars) significantly raised the pH of WCS417 growing in seedling exudates. (C) Arginine increased the pH of the Δ*pqqF* mutant and the Δ*argF*Δ*pqqF* double mutant, suggesting that arginine pathway is required for rhizosphere acidification. All the experiments were independently repeated at least 3 times. Statistics were calculated by using one-way ANOVA and Tukey’s HSD. Error bars represent mean +/− SD and letters indicate differences at p<0.05.

To test the hypothesis that accumulation of glutamate and ammonia could result in an increase in rhizosphere pH, we added arginine, proline or glutamine to seedling exudates containing bacteria, which should result in an accumulation of their precursors, glutamate and ammonia [17]. We also tested methionine, leucine, tryptophan, serine, and histidine, which should not affect glutamate catabolism, as controls. To avoid confounding our findings with acidification through gluconic acid biosynthesis, we added arginine to the *P. simiae* WCS417 Δ*pqqF* mutant. We found that exogenous arginine, proline, glutamine, and glutamate, but not methionine, leucine, tryptophan, serine, or histidine, significantly raised the pH of the Δ*pqqF* mutant close to the level of mock seedling exudates (Figure 3B). Furthermore, we found this alkalization phenotype is even more dramatic in PAO1 as arginine, glutamine and glutamate raised the pH of seedling exudates inoculated with PAO1 *pqqF*::Tn5 to around 8.0 (Figure S3). These data indicate that similar to pathogenic fungi, exogenous arginine, proline, glutamine, or glutamate likely results in rhizosphere alkalization through ammonia accumulation.

To test genetically whether a loss of arginine, proline, glutamine, or glutamate, but not other amino acids, could specifically result in increased pH of seedling exudates, we selected 6 mutants that have insertions in the genes that are required for amino acid biosynthesis from the *P. aeruginosa* PAO1 transposon insertion library (Table S3). We found that *proA*::Tn5 (deficient in proline biosynthesis) and *gltB*::Tn5 (deficient in glutamine biosynthesis) neither acidify seedling exudates nor suppress PTI (Figure S2A, B) consistent with glutamate being required for rhizosphere acidification. Two additional amino acid insertions, in *metZ*::Tn5 (deficient in methionine biosynthesis) and *leuC*::Tn5 (deficient in leucine biosynthesis) also resulted in a loss of acidification of seedling exudates and immunity suppression of seedlings suggesting that leucine and methionine may be limiting for bacterial growth in the rhizosphere (Figure S2A, B). In addition, *serA*::Tn5 (deficient in serine biosynthesis) suppressed host immunity and had decreased pH of seedling exudates indicating that the rhizosphere may contain enough serine to support bacterial growth (Figure S2A, B). Interestingly a *hisB*::Tn5 mutant (deficient in histidine biosynthesis) can also acidify seedling exudates but induced immunity on its own further indicating that low pH alone is not sufficient for immunity suppression (Figure S2A, B). Collectively these data confirm that the rhizosphere is deficient in glutamate and downstream amino acids and so bacteria must actively synthesize these in the rhizosphere.

Previous reports have shown that amino acid auxotrophs, including an *argF* transposon insertion mutant in WCS417, exhibit enhanced fitness in the *Arabidopsis* rhizosphere [18]. This suggests there is already sufficient arginine and other amino acids in the rhizosphere to support bacterial growth, and so *argF* should not be active. However, this contradicts our findings that arginine biosynthesis is required for pH regulation and immunity suppression. As a result, we tested whether arginine was limiting in our system by testing whether the Δ*argF* mutant had a growth defect in seedling exudates, and whether adding exogenous arginine could rescue growth. We found that indeed the *argF* mutant has a growth defect in seedling exudates suggesting arginine may be limiting for growth (Figure S4A). We found that while supplying exogenous amino acids had no effect on the ability of wildtype *P. simiae* WCS417 to suppress flg22-triggered *CYP71A12_pro_:GUS* expression, it restored the ability of the *argF* mutant to suppress immunity and reduced pH (Figure S4B). These data suggest that arginine levels in the rhizosphere may be limiting for bacterial growth, and that the lack of immunity suppression and acidification by the Δ*argF* single mutant may be due to its inability to grow.

Because *P. simiae* WCS417 produces large amounts of gluconic acid, and we only see an increase in pH with application of exogenous amino acids in the Δ*pqqF* mutant (Figure 3B), we hypothesized that *argF* might be necessary for lowering pH, but that gluconic acid might mask the effect of *argF*. If this is the case, we would expect that exogenous arginine would restore growth, but not lower the pH of a double Δ*pqqF*Δ*argF* mutant growing in seedling exudates. In contrast, if *argF* only contributes to growth, addition of exogenous arginine should restore pH of the Δ*pqqF*Δ*argF* mutant to the level of the single Δ*pqqF* mutant. Indeed, we found that the double mutant with arginine still had significantly higher rhizosphere pH than the Δ*pqqF* mutant alone (Figure 3C), but that exogenous arginine rescued the ability of a Δ*pqqF*Δ*argF* double mutant to suppress immunity (Figure S4). These data indicate that *argF* is required for both growth and regulation of pH in the rhizosphere. These data also indicates that while low pH is sufficient to supress immunity, it is not necessary in WCS417 suggesting that there is a second, pH-independent mechanism of immunity suppression by *P. simiae* WCS417. Finally, they indicate that gluconic acid and amino acid biosynthesis work through separate partially redundant pathways to regulate rhizosphere pH, and that the final pH may be determined by the balance of glucose and amino acid availability in the rhizosphere.

To test our prediction that rhizosphere alkalization is due to accumulation of ammonia from inhibiting the arginine biosynthesis pathway, we measured the ammonium concentration in seedling exudates. In contrast to our prediction that addition of arginine, glutamine or glutamate would increase ammonium, we found that the Δ*pqqF* mutant consumes more ammonium than wild type bacteria in the presence of these amino acids (Figure 4A). This suggests that when arginine biosynthesis is inhibited, ammonia is converted to a distinct compound that contributes to rhizosphere alkalization. Since spermidine is an alkaline polyamine compound that is derived from ornithine, a precursor of arginine, we wondered whether spermidine accumulation could underlie the increased pH in the *pqqF-deficient* mutants. We found that addition of exogenous ornithine or putrescine had no effect on the pH of WCS417 in seedling exudates and ornithine but not putrescine resulted in a slight increase in the pH of the Δ*pqqF* mutant (Figure 3B). Similarly, we observed no effect of ornithine addition to wildtype PAO1 or the Δ*pqqF* mutant (Figure S3). These data indicate that ammonia conversion to ornithine and spermidine are unlikely to explain the increase in pH.

**Figure 4.**
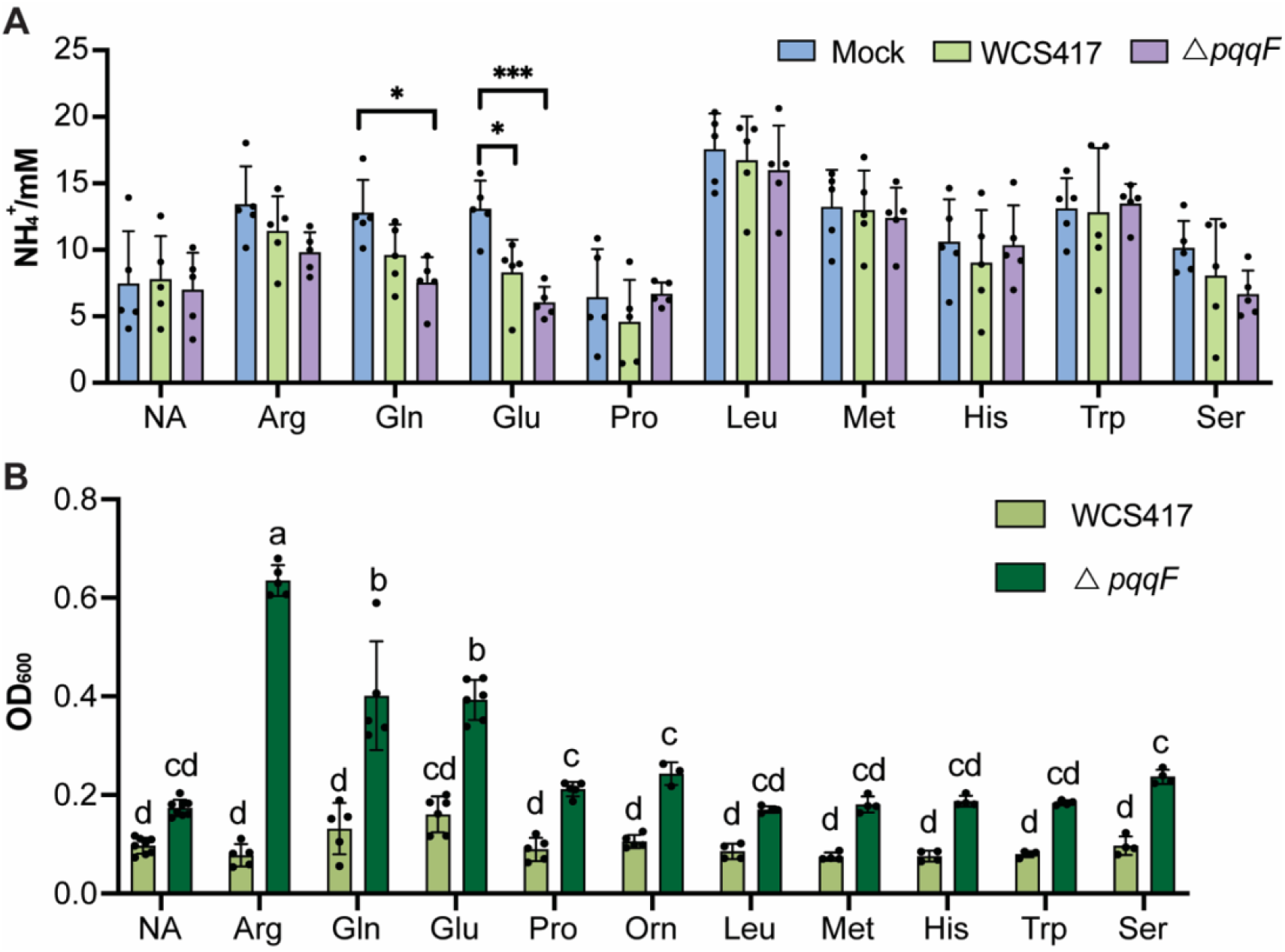
Alkalization correlate with bacteria growth but was not due to ammonia accumulation. (A) Ammonium concentration does not vary between the wildtype WCS417 and the Δ*pqqF* mutant. (B) Blocking glutamate biosynthesis by addition of exogenous arginine, glutamine, and glutamate results in bacteria overgrowth of the WCS417 Δ*pqqF*. All the experiments were independently repeated at least 3 times. Statistics were calculated by using one-way ANOVA and Tukey’s HSD. Error bars represent mean +/− SD and * indicate differences at p<0.05, *** indicate differences at p<0.01.

As fungal alkalization through glutamate secretion is accompanied by increased fungal growth and virulence [19], we tested whether an increase in pH might result in bacterial overgrowth. We observed that arginine, glutamine, and glutamate-mediated alkalization were accompanied by dramatic bacteria overgrowth of the both the WCS417 and PAO1 Δ*pqqF* mutants in both (Figure 4B, Figure S5). These data indicate that pH correlates with bacteria growth and that by limiting certain amino acids, plants can control the rhizosphere pH and bacterial growth.

### Acidification-mediated immunity suppression is peptide-specific in roots

Our data show that *Pseudomonas* have multiple, partially redundant mechanisms to acidify the rhizosphere indicating that maintenance of correct pH may be critical to establishing plant immune homeostasis. Previously, we found that acidification to pH 3.7 partially blocked flg22-mediated induction of the *MYB51_pro_:GUS* reporter gene, but had no effect on the expression of SA-triggered *NPR1_pro_:GUS* and *MYB72_pro_:GUS* triggered by WCS417 or WCS358 at pH 3.7 in roots, indicating that acidification can only impair a section of plant innate immunity [8]. The flg22 peptide is recognized by the extracellular leucine-rich repeat (LRR) domain of FLS2, and upon binding, induces association of FLS2 with the coreceptor BRASSINOSTEROID INSENSITIVE 1-associated kinase 1 (BAK1), activating a mitogen-activated protein kinases (MAPKs) cascade that activates defense genes expression [20,21]. Importantly, flg22-FLS2 recognition immediately induces extracellular alkalization and such binding requires an optimum pH between pH 5 and 6 [22]. As a result, we hypothesized that acidification likely impedes flg22-FLS2 interaction or FLS2-BAK1 ligand binding rather than intracellular MAPK activation or defense gene expression.

If acidification blocks flg22-FLS2 binding, then other MAMP-receptor interactions might not be affected if they have different binding affinities under low pH. We tested the effect of acidification on the damage-associated molecular pattern *At*pep1, which is BAK1-dependent [23,24] and chitin, a sugar polymer from fungal cell walls, which is BAK1-independent in *Arabidopsis* [25,26]. All three MAMPs flg22, *At*pep1 and chitin share a common MAPK cascade [27–29]. We tested whether acidification can also suppress chitin- and *At*pep1-triggered immunity using the MAMPs marker gene reporters *CYP71A12_pro_:GUS*. We found that at pH 3.7, *At*pep1-but not chitin-triggered expression of the *CYP71A12_pro_:GUS* reporter was abolished (**Figure 5A**). These data indicate that not all MAMP perception is inhibited suggesting that acidification likely interferences with receptor binding and not MAPK signaling.

**Figure 5.**
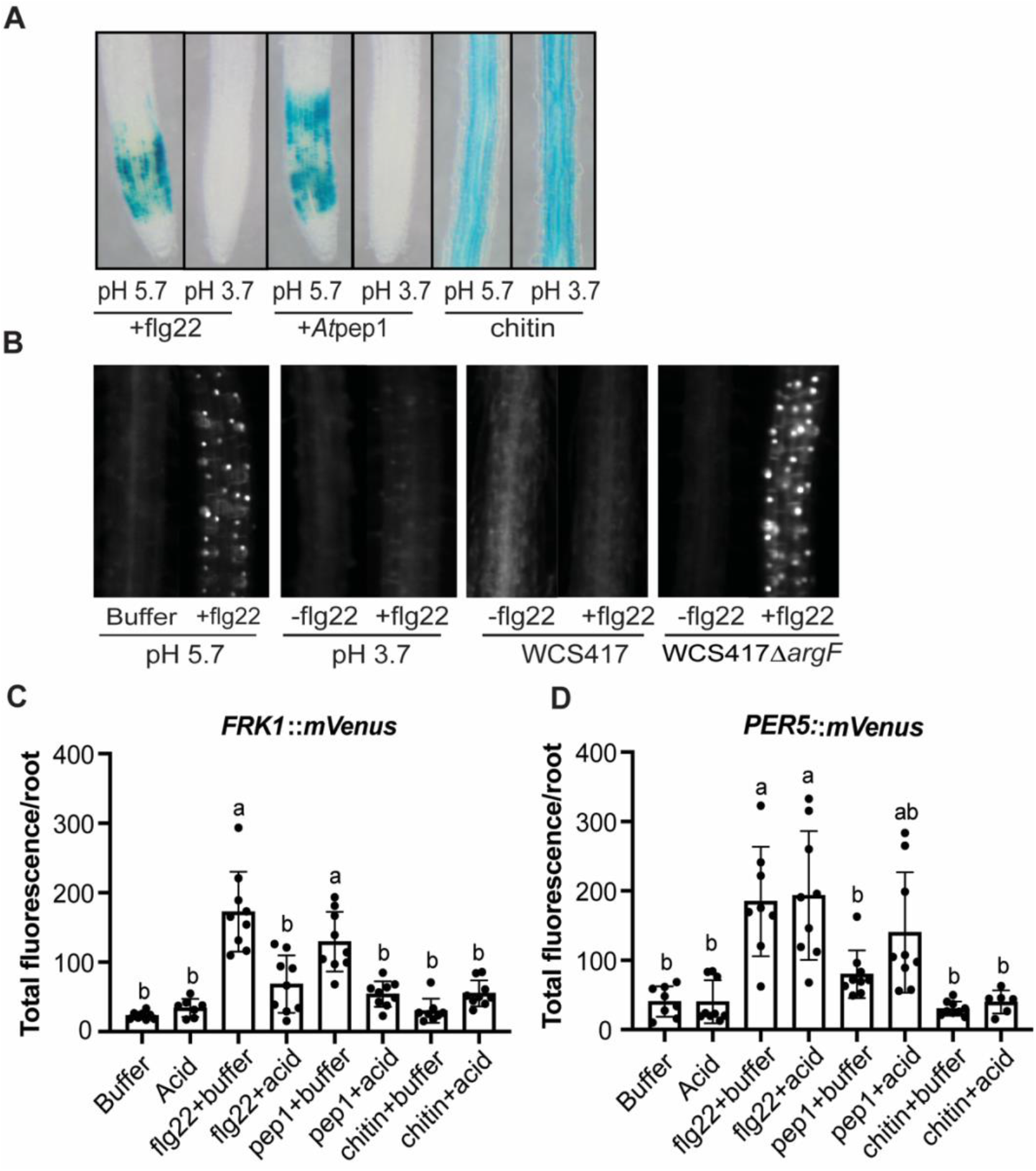
Acidification only suppresses a sector of plant innate immunity. (A) Acidification blocks flg22-but not Atpep1- or chitin-triggered *CYP71A12_pro_:GUS* expression. (B) Acidification and WCS417, but not the WCS417 Δ*argF* mutant can block flg22-induced *FRK1::mVenus* fluorescent reporter expression. (C-D) Acidification blocks flg22-induced *FRK1::mVenus* expression but not *PER5::mVenus* expression. All the experiments were independently repeated 3 times. Statistics were calculated by using one-way ANOVA and Tukey’s HSD. Error bars represent mean +/− SD and letters indicate differences at p<0.05.

We tested whether all PTI-induced gene expression was blocked by using additional PTI-inducible reporters. We used *PER5* (At1g14550) which is a peroxidase [30], and FRK1 (AT2G19190) which encodes a LRR receptor kinases [20]. Both genes are flg22 and *At*pep1-inducible in root [31–33]. We found that both flg22- and *At*pep1-induced *FRK1*::mVenus expression was significantly suppressed by low pH (Figure 5B, C). In contrast, lower pH did not affect flg22-induced *PER5::mVenus* expression or *At*pep1-induced *PER5::mVenus* expression (Figure 5D). These results suggest that acidification can only affect a sector of PTI and rhizosphere pH may be critical to determine the baseline setting on the plant immune thermostat.

## Discussion

It is intriguing that plants allow colonization of commensals but restrict pathogens through the plant immune system [34–37]. It is possible that plant immune system can distinguish commensals and pathogens, or that different bacteria possess strategies to avoid or suppress plant immunity [38,39]. Not only do taxonomically diverse root-associated commensals not induce immune responses, but they can elude or suppress host immunity [8,9,34,40]. Activation of the plant immune system reduces rhizosphere colonization and further shifts the microbiota community whereas suppression of host immunity enhances microbe colonization [7,34,41]. Therefore, rhizosphere-associated microbes must manipulate the plant immune system to promote their own fitness.

Here we report a forward genetic screen that identified a bacterial gene ornithine carbamoyltransferase *argF* from *Pseudomonas simiae* WCS417 that is required for host immunity suppression, colonization, and acidification. The Δ*argF* mutant is auxotrophic and without arginine and exogenous arginine restored Δ*argF*-mediated host immunity suppression, colonization, and acidification to wildtype levels. We tested other amino acid auxotrophs and found that the majority of them cannot acidify or suppress host immunity. This indicates that amino acid biosynthesis plays an important role in rhizosphere colonization. This is not the first time that amino acid biosynthesis has been shown to be necessary for root colonization [7,42].

Interestingly, a previous TnSeq screen found that amino acid auxotrophs, including insertions in *argF* in *P. simiae* WCS417 exhibited enhanced fitness in the *Arabidopsis* rhizosphere, which is the opposite of what we found [18]. We suspect the difference between these findings may be because they tested a community of transposon insertion mutants in a TnSeq screen, where the presence of other mutants could potentially provide amino acids in trans to auxotrophs. In fact, metabolic exchange, including amino acid cross-feeding among microbes is characteristic and reciprocal in a microbial community [43–45]. Our data suggest that the rhizosphere may be limiting in many amino acids, and that by synthesizing certain amino acids, bacteria will alter the rhizosphere pH and affect plant immune homeostasis.

To disentangle the role of amino acid biosynthesis in rhizosphere acidification from their role in growth, we supplied different amino acids to *Pseudomonas pqqF* mutants, which cannot produce gluconic acid but retains some rhizosphere acidification (Figure 2B). We found that only arginine, proline, glutamine, and glutamate caused rhizosphere alkalization and bacterial overgrowth in the *pqqF*-deficient mutants. We speculate that the alkalization was likely due to the accumulation of ammonia as this is a common strategy employed by fungal pathogens to promote host infection [15,16]. However, quantification of ammonium shows that ammonium levels do not explain the pH variation between the wildtype and the *pqqF*-deficient mutants. Thus, there might be a novel mechanism for glutamate pathway-mediated rhizosphere alkalization. In addition, arginine, glutamine and glutamate-mediated alkalization accompanies overgrowth of *pqqF-deficient* mutants, which is reminiscent of alkalization-mediated invasive growth in fungi [19]. This indicates that maintaining the balance of rhizosphere pH via glutamate and gluconic pathways is essential for bacterial growth regulation. Lastly, we noticed that providing serine also increased the pH of the WCS417 Δ*pqqF* mutant (Figure 3B), indicating that there might be another pathway or other compounds that contributes to rhizosphere acidification.

We found that while rhizosphere acidification is sufficient to suppress immunity, it is not necessary for immunity suppression in *P. simiae* WCS417 indicating there is a distinct mechanism of immunity suppression. The Δ*pqqF* mutant and the Δ*pqqF*Δ*argF* mutant supplemented with exogenous arginine caused an increased in rhizosphere pH, but still suppressed immunity. This indicates that there are additional pH-independent mechanisms for host immunity manipulation in *P. simiae* WCS417.

Rhizosphere acidification seems to be a general characteristic of many root-associated microbes [8,41]. Thus, rhizosphere acidification could be a conserved evolutionary trait of root-associated microbes. However, suppression of host immunity may also open the window for pathogens. In the present study, we found that acidification can only dampen a sector of immunity, indicating that plant must also evolve novel mechanisms to counteract acidification-mediated immunity suppression which may act as a selection force for microbial colonization.

Collectively, host immunity suppression is crucial for host colonization of both commensals and pathogens. Crosstalk between the microbes and the plant immune system is an ongoing process. Our results highlighted that apart from serving as colonization factors and nutrient, bacterial amino acid biosynthesis plays a novel dual role in the rhizosphere acidification and host immunity suppression (Figure 6). As acidification quenches only a sector of host immunity, it is clear that more mechanisms of host immunity suppression still have yet to be discovered.

**Figure 6.**
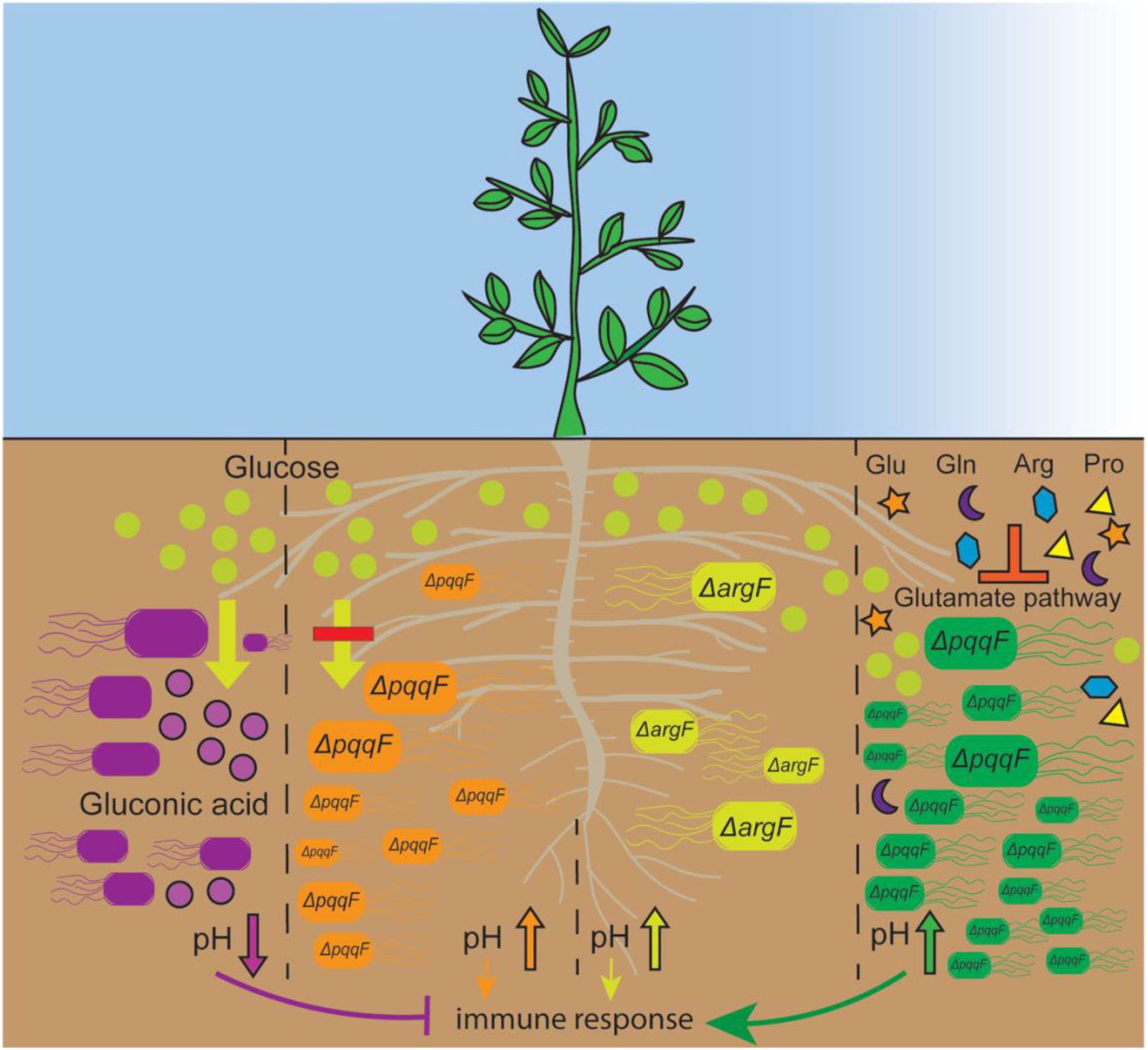
Model of bacteria-mediated rhizosphere acidification and acidification-mediated plant innate immunity suppression. Bacteria convert plant-derived glucose to gluconic acid to acidify the rhizosphere and suppress plant innate immunity. Abolishing *pqqF* in bacteria (orange bacteria) results in increased pH, increased bacteria growth, and in some cases, inability to suppress plant immunity. Bacteria actively synthesize certain amino acids (i.e., glutamine, glutamate, arginine, and proline) for survival since amino acid auxotrophs (i.e., Δ*argF;* yellow) exhibit rhizosphere growth defects, indicating that plants may restrict amino acids. When plants provide abundant glutamine, glutamate, arginine, or proline, which would block the glutamate biosynthesis pathway, it results in rhizosphere alkalization and significant bacterial overgrowth of *pqqF-deficient* mutants (green). Overall, this work suggests that maintaining an amino acid-glucose balance is crucial for regulating plant rhizosphere microbiome assembly.

## Materials and Methods

### Plant materials and growth conditions

We used *Arabidopsis thaliana* wild type Col-0, *CYP71A12_pro_:GUS* reporter [10] and *FRK1::mVenus, PER5::mVenus* [32] in our study. All the plant material used in our study were grown in the same condition, which was in a climate-controlled growth room at 21 °C, 16h light/ 8h dark cycle with light intensity of 100 μM. Plants were grown in the ½ X Murashige and Skoog (MS) media with 1% MES [2-(N-morpholino) ethanesulfonic acid] with 0.5% sucrose, adjusted by KOH to a pH of 5.7 [10].

### Bacterial strains and growth condition

Strains that were used in this study were listed in the Table S2. *Pseudomonas simiae* WCS417 and mutants were cultured overnight in LB or King’s B at 28°C with shaking at 180rpm. *Pseudomonas aeruginosa* PAO1 was cultured overnight in LB at 37°C with shaking at 180rpm. PAO1 transposon insertion mutants were obtained from the two-allele PAO1 transposon insertion library [12]. Wildtype PAO1 and the transposon insertion mutants used in this study were cultured in LB with 25μg/mL tetracycline at 37°C. *Escherichia coli* were cultured in 37°C with 15μg/mL or 100 μg/mL gentamycin depending on the experiment.

### ß-glucosidase (GUS) histochemical assays

The reporter lines *CYP71A12_pro_:GUS* in the *Arabidopsis* Col-0 genetic background contain the indicated promoter driving the expression of the ß-glucosidase reporter gene [10].

Seeds were grown in 48-well plates for one week after surface sterilization and were grown in the condition described above. Each well contained 300 μL ½x MS media and 0.5% sucrose. On day 8, the media was replaced by fresh ½x MS with 0.5% sucrose media. Bacteria were grown overnight in LB, washed in 10 mM MgSO_4_, and serially diluted to an OD_600_ of 0.02 in 10mM MgSO_4_. On day 9, 30 μl bacteria were added to each well (final OD_600_ of 0.002) and the plates were returned to the growth room for at least 18 hours before adding flg22. On day 10, flg22 was added to a final concentration of 500 nm and the media was replaced with GUS staining solution after 4.5 hours of incubation. The GUS staining solution was made fresh at a final concentration of 0.5 M sodium phosphate buffer (pH 7), 0.5M EDTA, 50mM potassium ferricyanide, 50mM potassium ferrocyanide, 50mM X-Gluc (5-bromo-4-chloro-3-indolyl-beta-D-glucuronic acid) and 10 μL triton X-100. Plates were then incubated at 37°C without light until the control roots treated with flg22 developed a visible blue color (approximately 3~4 hours). Finally, to clear the tissue, the GUS stain was replaced with 95% ethanol then washed with water afterwards. Images were taken with a Macro Zoom Fluorescence Microscope MVX10.

### Bacterial growth curves

Overnight cultures of bacteria were grown in LB, then were serially diluted to an OD_600_ of 0.2 in 10mM MgSO_4_ for growth curves. Growth curves were performed by adding 10 μL of the diluted culture to 90 μL rich media (LB), minimal media (M9 salts supplemented with 30mM succinate or 30mM glutamine), or seedling exudates (M9 salts supplemented with succinate, with or without 1mM arginine). Bacteria growth was quantified by measuring OD_600_ on a Versamax plate reader (Molecular Devices). Data presented in this study represent the average of three biological replicates.

### WCS417 EMS mutant library construction and screening

*Pseudomonas simiae* WCS417 (rif resistant variant) was mutagenized by spinning down and washing an overnight culture and exposing it to 1, 2 or 4% EMS for 1 hour. Mutagenized cells were plated on King’s B with 50 μg/mL streptomycin, and it was found that after treatment with 4% EMS there was ~100-fold increase in the number of resistant cells relative to the parental strain, so these cells were used for library construction. For library construction, mutagenized cells were plated on LB + rifampicin 50 μg/mL and individual colonies were picked into wells of 96-well deep well plates in LB media. Each plate contained 92 EMS mutants, and 4 wells containing the parental strain as positive controls. After overnight growth, 75 μl of LB containing library bacteria were pipetted into a fresh 96-well plate and 25 μL of 80% glycerol was added. The library was stored at −80°C.

To screen the library, seedlings were grown in 96-well plates in MS media as described above. Three seedlings were grown in each well containing 100 μL MS media. The library was stamped onto rectangular plates containing solid LB media, and then sub-cultured into 96-well deep bottom plates containing LB. After overnight growth, the OD_600_ of 4 independent wells was measured, and the average was taken. This was used to calculate an approximate dilution factor to dilute all 96 wells to a final OD_600_ = 0.05. 10 μl of the diluted culture was added to each well, containing 90 μL MS, for a final average bacterial concentration of 0.005. The screen was repeated in duplicate and candidates that failed to suppress immunity in both replicates were retested.

### Mapping WCS417 EMS mutations

To map WCS417 EMS mutations, we sequenced the genomes of the parental line used to make the library as well as the genomes of each individual mutant. Genomic DNA was extracted from WCS417 and the 10E10 EMS mutant with the Puregene Core Kit (Qiagen). A PE150 short insert library was prepared and sequenced on an Illumina HiSeq 2500 (Novogene). After adapter trimming with Cutadapt [46], reads were aligned to the WCS417 genome using the Bowtie2 [47] aligner with default parameters. Variant calling was performed with BCFtools [48]. Low quality SNPs with a quality score under 20 were filtered out and SNPs found in the parent strain were discarded from consideration.

### Bacteria mutant complementation

Complementation of the 10E10 mutant was performed by PCR amplifying the coding sequence and native promoter of *argF* in WCS417. The PCR product containing HindIII and BamHI restriction sites was ligated to the plasmid pBBR1MSC5 and the ligation product was transformed into the competent cell *E.coli* DH5α and plated on gentamycin 100μg/mL plates for selection of positive colonies. Positive colonies of *proargF*:pBBR1MSC5 construct were confirmed by colony PCR and further confirmed by Sanger sequencing. The confirmed pBBR1MCS-5::Pro_*argF*_-*argF* construct was then transformed into the 10E10 mutant.

### Generation of deletion mutants in WCS417

Clean deletion of *argF* or *pqqF* in WCS417 were made using a double-recombination method in Gram-negative bacteria using counter selection with *sacB* [49]. Two sets of primers were designed to amplify the 500bp flanking region upstream and downstream of *argF* or *pqqF*. Primer 1 with the restriction enzymes site HinIII and primer 4 with the restriction enzyme site BamHI are the left and right primers of the upstream and downstream flanking region of the target gene, respectively. Primer 2 and primer 3 are the right and left primers of the upstream and downstream flanking region of the target gene, respectively. Both primer 2 and primer 3 consistent of 15bp of region upstream and 15bp of region downstream of the target gene. Thus, primer pairs primer1 and primer2, and primer3 and primer4 were used to amplify the 500bp regions upstream and downstream of the target gene, respectively. Overlap PCR was performed with the upstream and downstream PCR products, which were digested with HindIII and BamHI and ligated to the pEXG2 suicide vector containing *sacB* [50], then transformed into *E.coli* DH5α. The positive colonies were selected on LB plates with gentamycin 15 μg /mL, then confirmed by colony PCR. The deletion constructs for *argF* or *pqqF* were further confirmed by Sanger sequencing. The confirmed *argF* or *pqqF* deletion constructs were then transformed into the competent SM10λ cells and was selected on LB plates with gentamycin 15ug/mL. Conjugation of the SM10λ containing the deletion construct and WCS417 was performed and the transconjugants were selected on plates containing nalidixic acid 15ug/mL and gentamycin 100μg /mL. Positive colonies were restreaked again and were cultured overnight in plain LB. Cell pellets were diluted to 10X and 100X and were each plated onto 10% sucrose plates and gentamycin 100μg/mL plates to select for the second recombination. The Δ*argF*Δ*pqqF* double mutant was made by conjugating the Δ*pqqF* mutant with the SM10λ strain containing the *argF*-pPEXG2 deletion construct and the selection was performed as described above.

### Seedling exudates

To generate seedling exudates, *Arabidopsis thaliana* Col-0 seeds were grown in half strength MS media containing 0.5% sucrose for 7 days as described above. The seedling exudates were collected from all the wells, immediately syringe filtered with a 0.22 μm filter and frozen at −20°C.

### Amino acid solutions

100 mM (100X) stock solutions of L-arginine, L-proline, L-glutamine, L-glutamate, L-ornithine, L-leucine, L-methionine, L-histidine, L-tryptophan, L-serine were made in water and filter sterilized using a 0.22 μm filter, before storing at 4°C.

### pH assay in seeding exudates

Bacteria were grown in LB overnight and were serially diluted to OD_600_ 0.02 in 10mM MgSO_4_. Bacteria were inoculated into 24-well plates containing seedling exudates to a final OD of 0.002 and the plates were incubated in a 28°C or 37°C incubator for 18 hours. Final concentration of 1mM amino acids (100X dilution of stocks) were added when required. The pH was measured using an ORION STAR A215 pH meter. 1mL of culture was directly taken from each well and the OD was measured by a spectrophotometer. Each experiment included three technical replicates and was independently repeated at least three times.

### Ammonium quantification

Ammonium concentration in the pH assay was measured by an Ammonia Assay Kit (Sigma, MAK310). The kit provides reagents that reacts with ammonia/ammonium ions, which produces fluorescence signals that are proportional to the ammonia concentration in the sample. 1 mL of each sample was taken from the pH assay and centrifuged for 5 minutes at 14,000 rpm. 10 μL of the supernatant was used for ammonia quantification. Ammonium quantification was performed following the manufacturer’s instructions. Plates were read by a Spectramax plate reader (λ_ex_ = 360/λ_em_ = 450nm).

### Fluorescence reporter imaging and quantification

*FRK1*::mVenus and *PER5*::mVenus seedlings were grown in 600 μL of ½x MS media with 0.5% sucrose and a pH of 5.7 in 24-well plates. On day 8, the media was replaced with 540 μL of fresh ½x MS with 0.5% sucrose with a pH 5.7. On day 9, MAMPs were added to a final concentrations of 500μM flg22, 100nM *At*pep1 or 0.1mg/mL chitin were added. For low pH condition, ½x MS with 0.5% sucrose with pH 3.7 were added along with the elicitors described above and incubated for 4.5 hours. Images were taken with a Macro Zoom Fluorescence Microscope MVX10 microscope.

## Acknowledgements

This work was supported by NSERC Discovery Grants NSERC-RGPIN-2016-04121 and NSERC-RGPIN-2021-03587 and Weston Seeding Food Innovation grants to C.H.H. Y.L. was supported by a Chinese Graduate Scholarship Council award and an NSERC CREATE-PRoTECT award. Early stages of this work were supported by NIH grant R37 GM48707 and NSF grants MCB-0519898 and IOS-0929226 awarded to Frederick M. Ausubel. We thank Dr. Bob Hancock for the *P. aeruginosa* PAO1 transposon insertion library. We thank Dr. Niko Geldner for providing the fluorescent reporters lines *FRK1::mVenus* and *PER5::mVenus*. We thank Dr. Yi Song and Dr. David Thoms for the training of the gnotobiotic plate assay and microscopic experiment, respectively. We thank Sarzana Hossain and Nicole R. Wang for insights and critical reading of the manuscript.

